# Conformational ensembles of the human intrinsically disordered proteome: Bridging chain compaction with function and sequence conservation

**DOI:** 10.1101/2023.05.08.539815

**Authors:** Giulio Tesei, Anna Ida Trolle, Nicolas Jonsson, Johannes Betz, Francesco Pesce, Kristoffer E. Johansson, Kresten Lindorff-Larsen

**Affiliations:** Structural Biology and NMR Laboratory & the Linderstrøm-Lang Centre for Protein Science, Department of Biology, University of Copenhagen, Copenhagen, Denmark

## Abstract

Intrinsically disordered proteins and regions (collectively IDRs) are pervasive across proteomes in all kingdoms of life, help shape biological functions, and are involved in numerous diseases. IDRs populate a diverse set of transiently formed structures, yet defy commonly held sequence-structure-function relationships. Recent developments in protein structure prediction have led to the ability to predict the three-dimensional structures of folded proteins at the proteome scale, and have enabled large-scale studies of structure-function relationships. In contrast, knowledge of the conformational properties of IDRs is scarce, in part because the sequences of disordered proteins are poorly conserved and because only few have been characterized experimentally. We have developed an efficient model to generate conformational ensembles of IDRs, and thereby to predict their conformational properties from sequence only. Here, we applied this model to simulate all IDRs of the human proteome. Examining conformational ensembles of 29,998 IDRs, we show how chain compaction is correlated with cellular function and localization, including in different types of biomolecular condensates. We train a model to predict compaction from sequence and use this to show conservation of structural properties across orthologs. Our results recapitulate observations from previous studies of individual protein systems, and enable us to study the relationship between sequence, conservation, conformational ensembles, biological function and disease variants at the proteome scale.

## Introduction

Intrinsically disordered proteins (IDPs) and regions (IDRs) within and between folded domains play important and highly diverse roles in determining and regulating biological processes. IDPs and IDRs—from hereon collectively IDRs—are highly dynamic molecules whose structures must be represented by conformational ensembles. This dynamics in turn complicates the interpretation of experimental data, and limits our ability to understand the relationship between protein sequences, conformational ensembles and biological functions of IDRs.

Recent developments in protein structure prediction have led to a transformative ability to predict structures of folded proteins, and to perform structure-based analyses of protein function at the proteome level [1]. When methods such as AlphaFold are applied to IDRs they, however, result in a single or few configurations of low confidence that do not generally represent the conformational heterogeneity and functional states of IDRs [2]. Indeed, the AlphaFold pLDDT confidence score is an accurate predictor of disorder, because the structures of disordered proteins are generally predicted with low confidence [1]. AlphaFold has, however, been shown to provide information about local regions of proteins that may fold conditionally [3, 4], and used to provide starting points for refinement of IDR ensembles against experimental data [5].

Structural studies of disordered proteins generally rely on combining molecular simulations with experimental data to correct for intrinsic biases in simulation methods [6]. Such studies have provided a wealth of insights for individual IDRs [7] and revealed sequence-ensemble relationships for individual types of proteins [8, 9], but do not provide information about the structural properties across the proteome. Our inability to predict structural properties of IDRs thus limits our understanding of the functional roles of IDRs and how evolution shapes them. As a supplement to these one-by-one studies of IDRs, we have recently taken a different approach where we used experimental data on more than 50 different proteins to learn the parameters in a molecular energy function to predict conformational properties of IDRs. By globally optimizing a transferable model, called CALVADOS, we can study the conformational ensemble of any IDR in the absence of experimental data, and have validated the model extensively [10, 11]. In developing CALVADOS we focused on predicting conformational properties such as long-range interactions and overall compaction, since these properties are known to be important for function [12] and the ability of IDRs to self-associate to form biomolecular condensates [9]; other methods already exist to predict local structuring or binding regions within disordered proteins from sequence alone [13].

We have here taken advantage of the speed and accuracy of CALVADOS to perform molecular simulations and generate conformational ensembles of all 29,998 IDRs from the human proteome. We analysed the conformational properties of the ensembles and related them to the biological function of the proteins. For example, we find that some functions such as the ability to bind RNA or the localization within nuclear speckles or spliceosomes are enriched in proteins that have compact IDRs, whereas components of chromatin and nucleosomes are enriched in expanded IDRs. Our large sequence-ensemble dataset reveals the sequence properties that determine conformational ensembles, including the interplay between overall hydrophobicity and charge patterning. We take advantage of this knowledge and our ensembles to train a model to predict protein compaction from sequence only, and use it to show that IDRs in human orthologs have conserved conformational properties. In a cellular context, many IDRs form interactions with other molecules, and chain compaction and charge properties of the monomer quantify key aspects of a protein’s ability to form both intra- and intermolecular interactions. Together, our work provides a new paradigm for how to link protein sequence, ensembles and function of disordered proteins, and sheds new light on the many biological roles of IDRs. Our database with conformational ensembles is freely available and will enable researchers to study the important and diverse roles of disordered proteins.

## Results and Discussion

### IDR selection and generation of conformational ensembles

We analysed AlphaFold2 models for 20,588 full-length human proteins and selected IDRs using low pLDDT confidence scores as predictor of disorder [1, 3, 4] (Fig. 1*A*), and obtained 29,998 sequences from 15,463 distinct proteins. The selected IDRs range in length between 30 and 500 residues (longer IDRs were divided into segments with a maximum of 500 residues), with a median value of 87 residues. Although pLDDT scores outperform many other disorder predictors, recent studies have highlighted that AlphaFold2 assigns high pLDDT scores to a fraction of known IDRs, possibly reflecting conditional folding upon binding or upon post-translational modifications [3, 4]. We therefore analysed sequence annotations in UniProt to calculate the fraction of residues within domains and structural motifs (*f*_domain_), and find that only a small fraction of the IDRs overlap substantially with annotated domains (Fig. S1).

**Figure 1.**
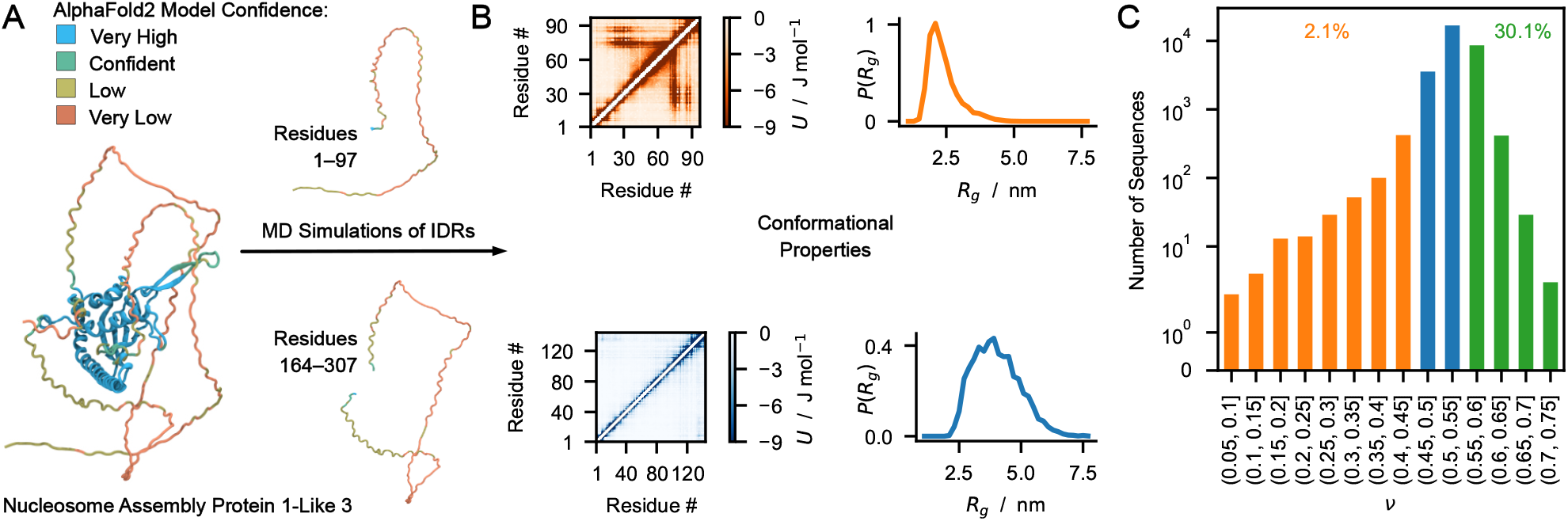
Schematic illustration of the approach used to obtain conformational properties for all the IDRs in the human proteome. (*A*) Selection of IDRs based on confidence of AlphaFold2 structural prediction, showing the selection of two IDRs from NP1L3 (UniProt ID: Q99457) as an example. (*B*) MD simulation of IDRs and calculation of conformational properties, showing interaction energy maps and distribution of the radius of gyration from the simulations of the two IDRs in NP1L3. (*C*) Distribution of the Flory scaling exponent, *ν* for 29,998 IDRs in the human proteome; note the logarithmic scale. A few IDRs have very small values of *ν* (0.3% have *ν* < 0.33); these IDRs display conformational properties that are not well described by a simple scaling law. We note that we do not find evidence for a significant enrichment of misclassified structured sequences in compact IDRs (Fig. S2).

We then performed MD simulations of all 29,998 sequences using the experimentally-parameterized and broadly validated residue-level CALVADOS model [10, 11] to generate conformational ensembles for each disordered sequence, and calculated distributions of conformational properties from these ensembles (Fig. 1*B*). The model was trained on experimental data reporting on single-chain conformational properties and was shown to predict radii of gyration of IDRs with mean relative error of 4% [11]. As measures of chain compaction that are independent of sequence length, we calculated the apparent Flory scaling exponent, *ν*, and the ratio of the mean-squared end-to-end distance and the mean-squared radius of gyration, 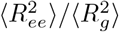; the accuracy of our model corresponds to an accuracy of predicting *ν* with an average error of ca. 0.01 for proteins of the lengths that we studied. For a so-called ‘ideal-chain’ polymer, the residue–residue, residue–solvent, and solvent–solvent interactions balance out and the polymer conformations are described by a Gaussian chain with *ν* = 0.5 and 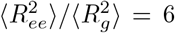 [14, 15]. For most of the simulated IDRs, we find conformational ensembles which approximate this ideal reference state. Only 2.1% of the IDRs are substantially more compact (*ν* ≤ 0.45) whereas 30.1% of the sequences adopt more extended conformations (*ν* > 0.55) (Fig. 1*C*). Although the relationship between 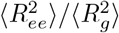 and *ν* is complex and sequence-dependent [15–18] (Fig. S3*A*), we obtain a similar distribution using 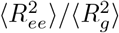 as a measurement of chain compaction (Fig. S3*B*). We also calculated other properties of the 29,998 conformational ensembles including interaction energy maps, asphericity (Δ) and prolateness (*S*). We find that both Δ and *S* are correlated with *ν* (Fig. S3*C*,*D*) so that the most compact ensembles are also most spherical and least prolate. We show 12 representative examples (Fig. S4) and provide access to all simulation trajectories and conformational properties in an online database. Generation of additional ensembles is possible via running CALVADOS using Google Colab (see Methods).

### Relating compaction with function and localization

The compaction of an IDR can play a determining role in protein function [12, 19, 20]. Chain compaction also indicates poor solvation, and may correlate with increased attractive chain-chain interactions and propensity to phase separate [9, 21, 22]. Although the coupling between conformational properties and phase behaviour may break for sequences with different net charges [23], IDRs of proteins that undergo homotypic phase separation tend to have lower *ν* values compared to IDRs of proteins that do not phase separate [24, 25]. For these reasons, we sought to investigate the association of the presence of IDRs in the most compact 5% (1,525 sequences with *ν* ≤ 0.477) with the biological function and cellular localization of the corresponding full-length proteins. Our gene ontology analysis shows a significant enrichment of compact IDRs in proteins that bind histones (OR = 2.8), RNA (odds ratio, OR = 2.0), chromatin (OR = 1.7), and DNA (OR = 1.4) (Fig. 2*A*). These data resonate with the strong association between proteins with compact IDRs and their localization to nuclear complexes and membraneless organelles, namely, the SWI/SNF chromatin-remodelling complex (OR = 3.0), the spliceosomal complex (OR = 2.9), nuclear speckles (OR = 2.1), the nucleolus (OR = 2.0), and nuclear bodies (OR = 1.7) (Fig. 2*B*). Conversely, the most expanded 5% of the IDRs (1,562 sequences with *ν* > 0.581) are significantly enriched in chromatin (OR = 25.8), in proteins that bind fibrillar collagen (OR = 18.8), in transmembrane proteins mediating the uptake of organic anions (OR = 5.9), and in chaperones (OR = 5.1) (Fig. 2).

**Figure 2.**
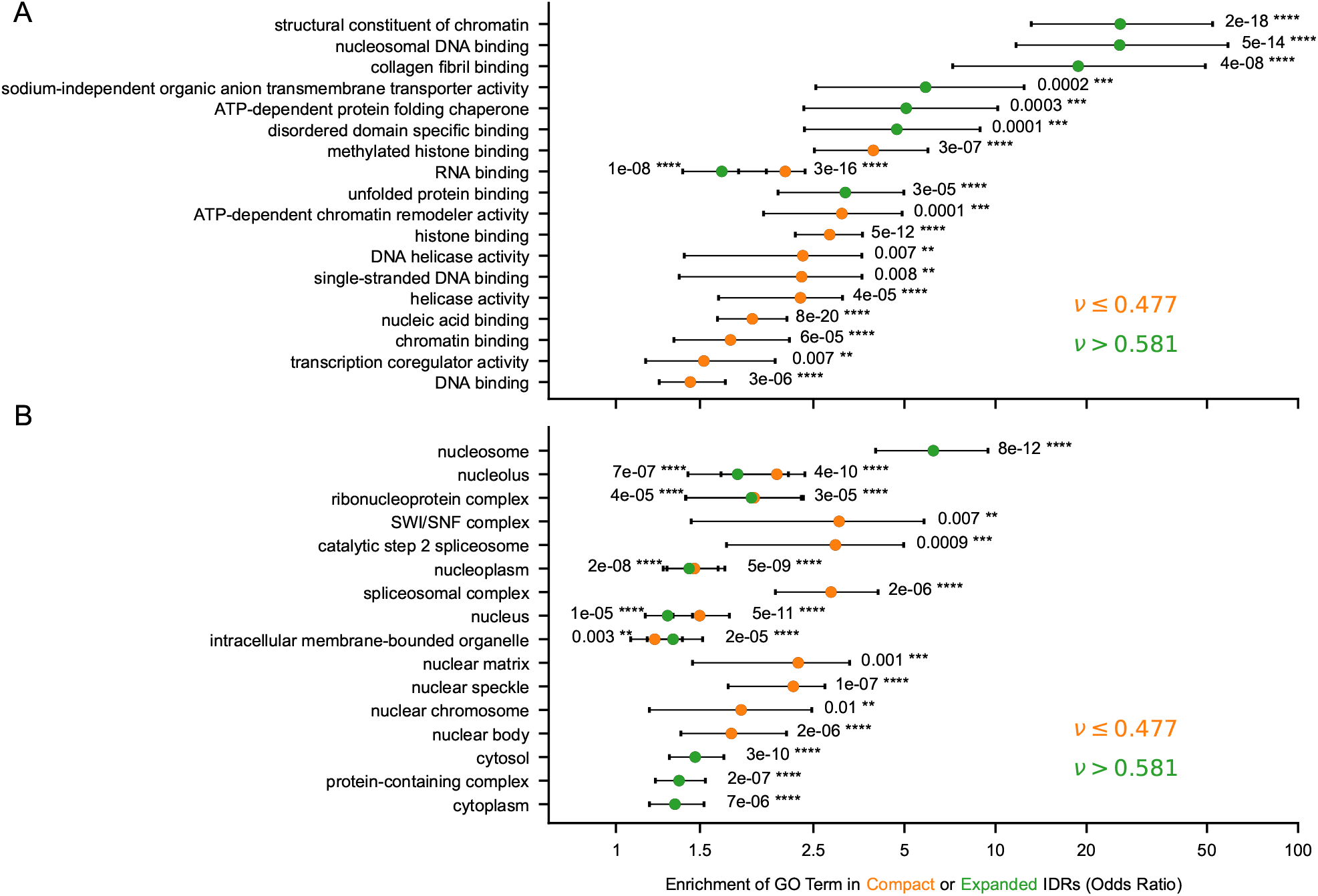
Gene ontology enrichment analysis of the most compact (*ν*≤ 0.477, orange) and most expanded (*ν* > 0.581, green) 5% of the IDRs. The strength of the association of compaction with (*A*) molecular functions and (*B*) cellular compartments is quantified by odds ratios. The statistical significance is estimated by adjusted *p*-values using Fisher’s exact test. Error bars show 95% confidence intervals.

We could ascribe a molecular function and cellular component annotations to 1,166 and 1,358 of the 1,525 IDRs with *ν* ≤ 0.477. Among these, we for example identified 27 previously analysed [26] activation domains of transcription factors, such as pancreas transcription factor 1 subunit alpha, forkhead box proteins O4 and O6, and the zinc finger proteins 473, 561, 644, and 777. Furthermore, as examples of compact IDRs of RNA-binding proteins, we find the well-characterized low-complexity domains hnRNPA1, hnRNPA2, and FUS, whose compaction correlates with their ability to form biomolecular condensates in the cell [9, 27–29]. Multivalent interactions between IDRs are one of the driving forces for the formation of membraneless organelles [30] and our gene ontology analysis suggests a strong association between IDR compaction and protein partitioning into nuclear condensates. We identified compact IDRs for 472 proteins that localize in intracellular membraneless organelles and for 232 proteins that are annotated to localize in nucleoli, nuclear speckles, nuclear bodies, and PML bodies. For example we predicted compact IDRs for nucleophosmin, a component of the nucleolus [31]; the nuclear speckle protein snRNP subunit U1-70k, which is a splicing factor and forms insoluble aggregates in Alzheimer’s disease [32, 33]; DAXX and ATRX, which are involved in chromatin remodeling and localize to PML bodies [34]; and CTR9, which is a positive regulator of RNA polymerase-II-mediated transcription [35].

### Insights into sequence–compaction relationships

While the positional sequence identity of IDRs is generally poorly conserved [36, 37], amino acid composition and patterning have been proven useful for predicting conformational and phase properties as well as for identifying functionally similar IDRs [9, 26, 33, 35, 38–46].

Segregation of charged residues into oppositely charged blocks has been identified as a key driver of chain compaction [38, 39, 41] and condensate formation [44,45, 47]. Conversely, high net charge per residue and uniform charge distribution result in expanded conformations [38, 39] and disfavor phase separation [10, 23, 48]. Recent studies have highlighted that IDRs with pronounced charge segregation are enriched in proteins involved in regulation of gene expression and mRNA processing. Specifically, for the IDRs of RNA polymerase II regulators, splicing factors, and polyadenylation factors, the extent of charge segregation has been shown to influence the selective partitioning of these proteins into transcription condensates [35], nuclear speckles [33], and nucleoli [46]. Moreover, charge blockiness has been identified as a conserved sequence feature [40] for example in homologous IDRs of the transcription elongation factors CTR9 and SPT6 [35].

Likewise, hydrophobic patterning modulates chain compaction and the propensity of IDRs to form condensates and avoid aggregation [9, 41]. Proteome-wide analyses have shown that the distributions of aromatic residues in prion-like IDRs is significantly more uniform than for random sequences [9, 49]. This finding suggests an evolutionary bias toward multivalent interactions between aromatic residues which underlie condensate formation, as opposed to stronger interactions between aromatic clusters which promote irreversible aggregation [9]. However, hydrophobic clusters have also been related to specific biological functions, as for the activation domains of transcription factors that bind coactivators. In these IDRs, clusters of aromatic and leucine residues are embedded in sequences rich in negatively charged residues [50], which may prevent chain collapse via electrostatic repulsion [26] and possibly degradation [51].

Inspired by these previous studies of individual protein families, we sought to examine the relationship between sequence and compaction across the entire set of human IDRs. For each sequence, we estimated the average residue stickiness, ⟨λ⟩ [11], as a measurement of the strength of attractive intra-chain interactions relative to solvent-protein interactions, and quantified the patterning of hydrophobic residues though the sequence hydropathy decorator (SHD) [41], using the amino-acid-specific stickiness parameters of the CALVADOS model. As metrics of charge segregation, we calculated the *κ* parameter [39] and the sequence charge decorator (SCD) [38]; *κ* and SCD capture different aspects of how uniform the charged residues are distributed along the polypeptide chain. More pronounced charge segregation is indicated by increasing *κ* values and decreasing SCD values. The *κ* parameter is normalized based on the composition of each sequence and ranges from 0 to 1 whereas SCD values correlate with the absolute net charge of the sequences. Therefore, *κ* and SCD are both influenced by sequence length and composition. Moreover, since *κ* is based on averages over windows of 5–6 residues [39], only SCD captures long-range interactions between blocks of charged residues along the linear sequence [48, 52].

We used the full set of IDR sequences to determine the sequence properties that affect chain compaction in human IDRs (Fig. 3). We find independent effects of SCD and *κ* (Fig. 3*A,B,G*); large negative and positive SCD values effectively distinguish chains with *ν* ≈ 0.5 from more compact and expanded IDRs, respectively (Fig. 3*A,E*). On the other hand, sequences with low or high values of *ν* are associated with larger *κ* values (Fig. 3*B,F*). With increasing compaction, sequences show increasing average stickiness, ⟨λ⟩, up to *ν* ∼ 0.5, after which ⟨*λ*⟩ plateaus and further decreases while the segregation of charged residues abruptly increases (Fig. 3*A*–*C*). These trends suggest that while hydrophobicity is used to drive some compaction, more extreme compaction is driven by electrostatic interactions between blocks of oppositely charged residues, in agreement with results from smaller sets of designed sequence variants [8, 39], and with the analysis of simulations of 16,885 synthetic sequences of equal chain length in an analysis performed in parallel with ours [53].

**Figure 3.**
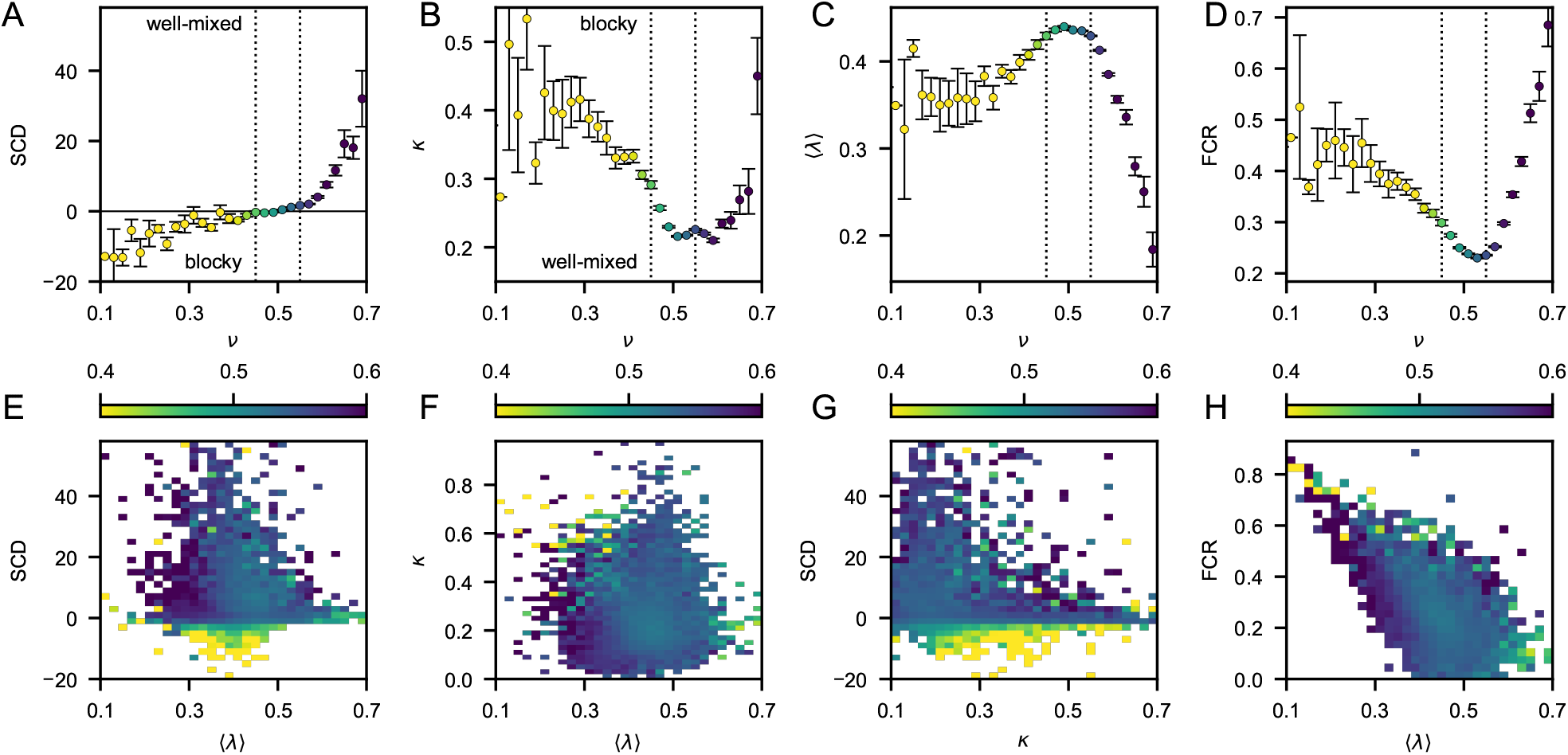
Sequence features that determine compaction. *A*–*D*: We show (*A*) sequence charge decoration, SCD, (*B*) charge segregation, *κ*, (*C*) average stickiness, ⟨λ⟩, and (*D*) fraction of charged residues, FCR as a function of the Flory scaling exponent, *ν. E*–*H*: Average *ν* as a function of (*E*) SCD and ⟨λ⟩, (*F*) *κ* and ⟨λ⟩, (*G*) SCD and *κ*, and (*H*) FCR and ⟨λ⟩.

We hypothesized that Nature evolved the most compact IDRs by exploiting charge segregation rather than increased hydrophobicity because sequences with large ⟨λ⟩ might lead to aggregation or degradation by the quality control apparatus [54]. To investigate this hypothesis, we used a predictor of quality control degradation signals, QCDPred, to estimate the propensities of the IDRs to be degraded [51]. The average QCDPred scores are low across the whole range of compaction, and more so for both compact and extended IDRs (Fig. S5*A*). We note that compact IDRs have on average considerably longer sequences than extended IDRs (Fig. S5*B*) and that the average QCDPred scores systematically decrease with increasing sequence length (Fig. S5*C*). This bias does therefore not explain the pronounced decrease in QCDPred score with decreasing *ν* below *ν* = 0.45 and the similar values of the QCDPred score for extremely compact and expanded IDRs, which have significantly different hydrophobic content. Instead, the excess of negatively charged residues with respect to arginine and lysine residues in compact IDRs (Fig. S6) may weaken the degradation signal, as suggested by recent experiments [55].

Whereas both compact and expanded IDRs are characterized by large fractions of charged residues (FCR) (Fig. 3*D*), expanded sequences are short, acidic, and depleted in arginine residues (Fig. S5*B* and S6). Conversely, compact IDRs are considerably longer, charge segregated, and richer in positively charged residues, although they have on average negative net charge per residue (Fig. S6*C*). Notably, IDRs in the most compact 5% have more arginine (⟨*f*_*R*_⟩ = 7.7 ± 0.1%) than lysine residues (⟨*f*_*K*_⟩ = 6.4 ± 0.1%). Because of the amphiphilic character of the guanidinium side chain [56], and its ability to favorably interact with aromatic residues [23], this imbalance may contribute to stabilize compact conformations as well as protein-protein and protein-nucleotide interactions [27, 57].

### Conservation of conformational and sequence features across homologs

Prompted by the observed relationships between sequence, compaction and biological function, we investigated the conservation of *ν* and sequence descriptors across a previously curated set of orthologs of human IDRs [58]. To estimate the scaling exponents of all the sequences, we trained a support vector regression (SVR) model against the *ν* values obtained from our simulations. Using permutation feature importance testing [59], we ranked the sequence descriptors based on their impact on model performance and selected the first five as input variables, namely, SHD > SCD > *κ* > FCR > ⟨*λ*⟩ (Fig. S7*A*). For a test set of 2,999 IDRs, the *ν* values predicted by the developed model have a Pearson correlation coefficient with the simulation data of 0.87 ± 0.02 and a mean absolute error of ca. 0.01 (Fig. 4*A*). Moreover, the SVR model accurately recapitulates the relationships between sequence features and *ν* obtained from simulations (Fig. S7*B*–*F*).

**Figure 4.**
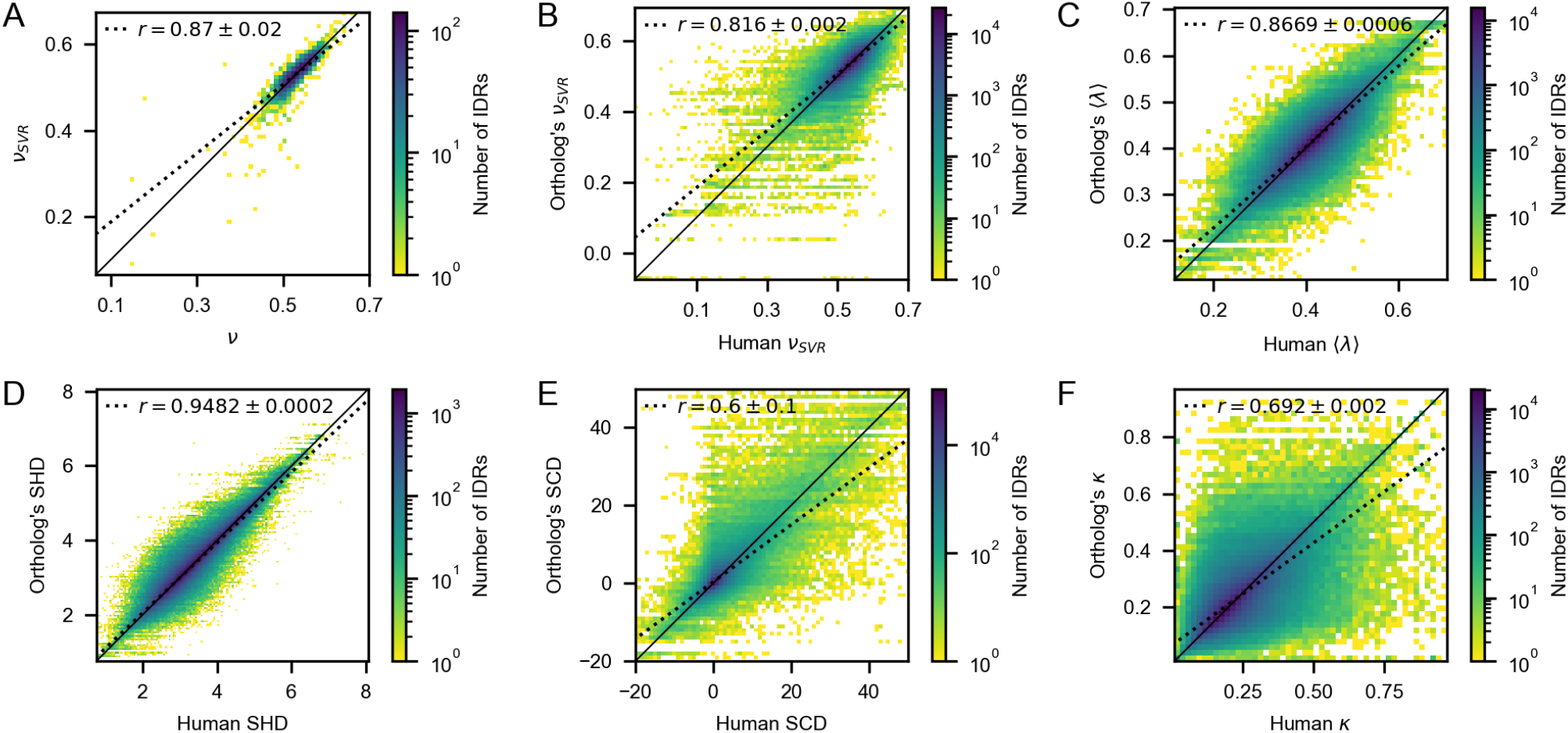
Conservation of conformational and sequence properties. (*A*) Correlation between *ν* values from simulations and *ν*_SVR_ values predicted using the SVR model for a held-out test set of 2,999 sequences from the set of 29,998 IDRs identified in this work. (*B*–*F*) Correlation of *ν*, ⟨*λ* ⟩, SHD, SCD, and *κ* between human IDRs and their orthologs from a set of 513,333 homologous IDRs, ranging in sequence length from 30 to 5,601 residues [58]. Dotted lines show linear fits to the data. *r* is the Pearson correlation coefficient with the error estimated as the standard deviation over 1,000 bootstraps.

Using the SVR model, we calculated *ν* for the set of the 507,059 homologous IDRs (15,140 human IDRs and their 491,919 orthologs) [58] and found a high correlation between the compaction of human sequences and their orthologs (Fig. 4*B*). Average chain stickiness and hydrophobic patterning are also highly correlated between human and orthologous sequences (Fig. 4*C,D*). These findings are in agreement with recent bioinformatics analyses on homologous prion-like domains of FUS and FET family proteins, which showed that sequence features governing chain compaction are evolutionarily constrained [9, 23, 49, 60].

Compared to ⟨*λ*⟩ and SHD, we observed a weaker correlation for both SCD and *κ* between human and homologous IDRs (Fig. 4*E,F*). Despite the importance of charge segregation in governing chain compaction, this finding may suggest that this feature is less evolutionarily conserved than hydrophobic patterning. This weaker correlation observed may, however, possibly be attributed to the effect of post-translational modifications on charge patterning [61, 62], which may differ across species and was not accessible for our analysis.

### Incidence of pathogenic variants and understudied genes

Given the observed evolutionary conservation of compaction, and its significant association with biological functions and cellular localization, we sought to investigate the potential relationship between IDR compaction and the incidence of pathogenic variants. We searched the ClinVar database [63] for missense variants in the human IDRs and found 2,977 pathogenic and 16,890 benign variants in 4,334 distinct proteins and 7,001 IDRs, which is less than a fourth of the IDRs in our set. Known pathogenic variants tend to be in regions with higher pLDDT scores than benign variants [64] (Fig. S8); this enrichment might be partially explained by the presence of a small fraction of folded regions within our set of IDRs (Fig. S8*B*), but may also reflect increased sensitivity in conserved regions of IDRs or those that may fold upon binding to their targets [3]. The sparsity of clinical data for IDRs could in part be ascribed to (i) the unknown functions of many IDRs and (ii) because computational methods to predict variant effects based on structure and sequence conservation are less applicable [13, 65]. Nonetheless, pathogenic variants in IDRs have been associated with many diseases [66], such as autism spectrum disorder, cancer, and neurodegenerative disorders [65]. Specifically, several disease-associated variants in IDRs have been shown to alter the formation, composition, and material properties of biomolecular condensates [67–71]. Our analysis shows that expanded chains have on average a lower number of pathogenic and benign variants than more compact chains (Fig. 5*A*). However, this trend can be attributed to the sequence length of the expanded IDRs, which are on average approximately three times shorter than compact IDRs (Fig. S5*B*), and we do not find evidence for an overall enrichment of pathogenic variants in more compact IDRs. We used our SVR model to predict the change in *ν* for the 2,977 pathogenic and 16,890 benign missense variants in ClinVar and find very similar distributions for the two classes suggesting that changes in compaction of IDRs is not a general disease mechanism for these variants (Fig. S9*A,B*). Similar results were obtained for 5,676 frameshift variants that have recently been described to lead to modified disordered C-terminal regions [70] (Fig. S9*C,D*).

**Figure 5.**
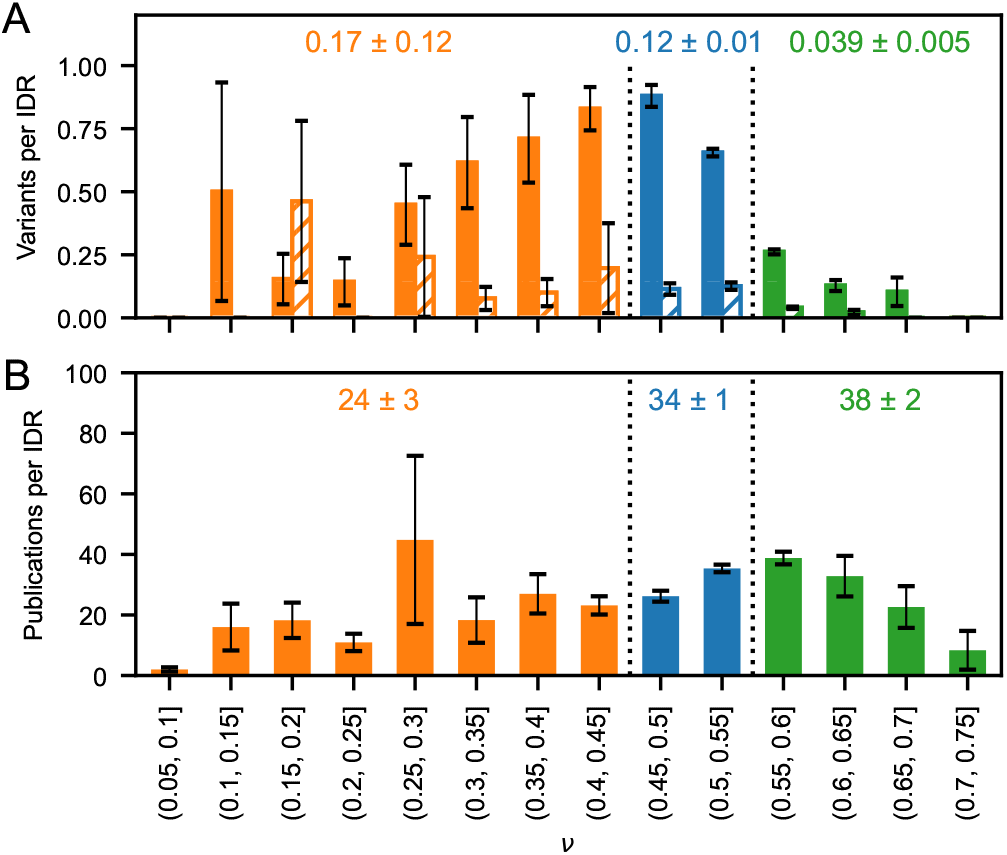
(*A*) Average number per IDR of benign (full bars) and pathogenic (hatched bars) ClinVar missense variants. (*B*) Average number of publications per IDR obtained from the “Find my Understudied Gene” tool [72]. Error bars show standard errors and the numbers refer to an average over the IDRs in the three groups of compaction.

The ability to study biological functions and assign functional status to missense variants depends on our knowledge of protein function, which varies substantially across the proteome [73]. We therefore asked whether certain IDR-containing proteins are less studied than others [72]. We calculated an average of 35 publications per IDR and observed an uneven distribution with respect to chain compaction, which suggests that proteins with the most compact IDRs may have received less research attention than those with more expanded IDRs (Fig. 5*B*), perhaps because they are longer and more difficult to characterize experimentally.

## Conclusions

IDRs are found in all kingdoms of life, and play essential roles in human biology and pathology. Our ability to study IDRs is, however, hampered by the fact that the relationship between sequence and structure is different from the more clearly defined patterns in folded proteins. This in turn makes it difficult to design experiments to test sequence-ensemble-function studies, and to understand the evolutionary processes that shape these. We have here performed a systematic analysis of the relationship between conformational ensembles and biological functions of human IDRs. Our conformational ensembles help rationalize previously observed relationships between sequence and function, and provides a unique dataset to continue to expand our knowledge of this understudied class of proteins, including training models to predict conformational properties [53, 74] or generate conformational ensembles [75]. In a cellular context, IDRs form interactions with other molecules, and our work provides a starting point for understanding both homo- and heterotypic interactions in IDR biology. Together with our and related methods [53] for predicting conformational properties from sequence and the mapping of biological observations onto our ensembles, we hope that our database of conformational ensembles presented in this work will encourage the investigation and the generation of hypotheses regarding the biological function, cellular localization, and variant effects of IDRs.

## Methods

### Selection of IDRs

Window-averaged AlphaFold2 pLDDT scores [1] were mapped to 20,215 (> 98%) sequences from the human proteome as defined in UniProt release 2021 4. Residues with ⟨pLDDT⟩ > 0.8, ⟨pLDDT⟩ < 0.7, and 0.7 ≤ ⟨pLDDT⟩ ≤ 0.8 were initially labeled as folded, disordered, and gap regions, respectively. Next, folded and disordered regions shorter than 10 residues were reclassified as gaps. Gap regions were relabeled as disordered if (i) flanked by disordered regions or (ii) terminal and preceded or followed by disordered regions. All other gap regions were instead relabeled as folded. IDRs smaller than 30 residues were discarded whereas IDRs larger than 500 residues were split into approximately equally long segments. This set of IDRs contain 37% of the residues in the UniProt proteome.

### Analysis of domain annotations

AlphaFold2 may in some cases assign high pLDDT scores to IDRs, possibly reflecting conditional folding upon binding or upon post-translational modifications [3, 4]. Moreover, the absence of cofactors or ions in AlphaFold2 predictions may lead to low pLDDT scores for domains such as zinc fingers, EF-hand motifs, and C2 domains [76]. To assess the extent of folded residues in the selected IDRs, we analysed domain annotations in UniProt; for each IDR we calculated the fraction of residues within annotated domains and structural motifs, *f*_domain_. Specifically, we obtained domain and zinc finger annotations for the selected IDRs from UniProt (uniprot.org/help/api) via application programming interface (API) calls (April 2023). A minimum sequence overlap of ten residues was used to evaluate the presence of the domains in the IDRs. We found that only 2.1% of the sequences have *f*_domain_ *>* 0.5 whereas 1.0% have *f*_domain_ *>* 0.9 (Fig. S1*A*). These putative folded sequences have generally short sequence length (Fig. S1*B*) and often include zinc or calcium binding sites, loops in protein kinase and myosin motor domains, and disulfide bonds in extracellular protein domains (Fig. S1*C*).

### Molecular simulations

Molecular dynamics simulations were conducted in the *NV T* ensemble at *T* = 37 ºC using the Langevin integrator with a time step of 10 fs and friction coefficient of 0.01 ps^−1^. The C*α*-based model was implemented using the CALVADOS 2 parameters and functional forms as previously detailed [11]. Single chains of *N* residues are simulated using HOOMD-blue v2.9.3 [77] in a cubic box of side length 0.38 × (*N* − 1) + 4 nm under periodic boundary conditions, starting from the fully extended linear conformation. The charge number of the first or last residue of the simulated sequences is increased or decreased by one unit when it coincides with the N-or C-terminus of the full-length protein, respectively. Histidine residues are modeled in the neutral form. Conformations are saved every Δ*t* ≈ 3 × *N* ^2^ fs if *N* > 150 and Δ*t* = 70 ps otherwise. Each chain is simulated for a simulation time of 1, 010 × Δ*t* and the initial 10 frames are discarded so as to obtain 1,000 weakly correlated conformations for each protein (Fig. S10*A*). The uncertainty of the averages of the radius of gyration, *R*_*g*_, and of the end-to-end distance, *R*_*ee*_, over the conformational ensembles is estimated as the standard error obtained from the blocking approach [78] implemented in the BLOCKING software (github.com/fpesceKU/BLOCKING). Apparent Flory scaling exponents, *ν*, are obtained by fitting 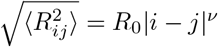 to root-mean-square residue-residue distances, 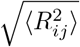, calculated for separations along the linear sequence, |*i* − *j*|, greater than 5. The sampling error on *ν* and 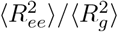 was estimated to be 0.01 and 0.1, respectively, from the standard deviation over five independent replicas averaged over a subset of 33 IDRs (Fig. S10*B,C*). Intra-chain distances were calculated from simulations trajectories using MDTraj [79]. The charge segregation parameter *κ* was calculated using localCIDER [80] or set to zero for sequences without charged residues.

We also provide easy access to generate conformational ensembles using CALVADOS simulations via a Notebook that for example can be run using Google colaboratory github.com/KULL-Centre/EnsembleLab. Typical simulation times range from ca. 5 min (a 71 ns-long simulation of a protein of 70 residues), 7 min (71 ns-long simulation of 140 residues), to 34 min for a 373 ns-long simulation of a protein with 351 residues.

### Gene-Ontology analysis

Gene Ontology (GO) terms for the selected IDRs were obtained directly from UniProt (uniprot.org/help/api) via API calls (January 2023). The basic version of the GO was downloaded in OBO format from the GO Consortium (purl.obolibrary.org/obo/go/go-basic.obo) (January 2023) and the Python package for complex network analysis NetworkX [81] was used to read the file as a directed acyclic graph. The graph has a hierarchical structure with three root nodes: “cellular component”, “biological process”, and “molecular function”. In this work, we analysed the “molecular function” and “cellular component” domains and discarded all child terms of the “biological process” domain. This filtering resulted in 23,916 and 27,129 IDRs with GO annotations of the corresponding full-length proteins pertaining to “molecular function” and “cellular component”, respectively. The numbers of compact IDRs (*ν* ≤ 0.477) with “molecular function” and “cellular component” annotations are 1,166 and 1,358 whereas the corresponding numbers for expanded IDRs (*ν* > 0.581) are 1,334 and 1,460. The GO terms were mapped onto parent terms occurring at least 10 times in proteins of IDRs with *ν* ≤ 0.477 or *ν* > 0.581. GO terms which are unrelated to any of the selected parent terms were labeled as “other”. The GO terms were divided into two bins based on the *ν* values of the IDRs and the thresholds *ν* ≤ 0.477 or *ν* >0.581. For each GO term, a 2 × 2 contingency table was populated with the numbers of occurrences of the GO term for IDRs with *ν* below or above the threshold. The strength of the association between chain compaction and GO terms was quantified as the odds ratios

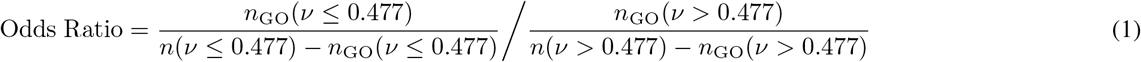

and

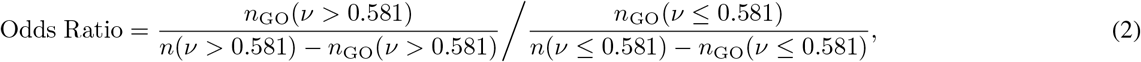

where *n*_GO_ and *n* denote the number of occurrences of the GO term and the total number of sequences in the *ν*-bin, respectively. Fisher’s exact test was used to estimate the significance of the association and the *p*-values were adjusted for multiple testing according to the Benjamini-Hochberg method [82].

### QCD calculations

We used QCDPred [51] to predict quality control degrons for all IDR sequences, and report the mean degron score across each IDR as representing its degron potential. Briefly, this method considers all possible 17-residues segments of an IDR and predicts a degron score based on the amino acid composition with hydrophobic residues acting to promote degradation and negatively charged residues to counteract degradation.

### Support vector regression model

A support vector regression model with a radial basis function kernel [83, 84] was developed to predict *ν* using as input features the sequence charge decoration, SCD [38]; the sequence hydropathy decoration, SHD [41]; the mean stickiness ⟨*λ*⟩; the *κ* parameter [39]; and the fraction of charged residues, FCR. The model was generated by randomly splitting the simulated sequences into training (30%), validation (60%), and test sets (10%). The sequence descriptors SCD, SHD, ⟨*λ*⟩, *κ*, and FCR were selected from a larger set which also included the net charge per residue, NCPR; and the fraction of aromatic, *f*_aro_; arginine, *f*_R_; lysine, *f*_K_; glutamate, *f*_E_; and aspartate residues, *f*_D_. The selection was based on their impact on model performance estimated using permutation feature importance testing [59]. Optimal hyperparameters (*ϵ* = 0.022, *C* = 51) were obtained using grid search with the Pearson correlation coefficient as a performance metric, leading to a correlation of 0.87 ± 0.02 and a mean absolute error of 0.012 on the test set. All model training and application, including permutation feature importance testing [59], was performed using scikit-learn [85]. We note that a similar model was recently described [74] using simulations with a different force field [41] that generally puts emphasis on sequence hydrophobicity and does not capture experiments as accurately as CALVADOS [10].

### ClinVar analysis

ClinVar data were obtained by parsing the ftp.ncbi.nlm.nih.gov/pub/clinvar/tab_delimited/variant_summary.txt.gz (May 2021) as described previously [86]. Namely, we included only entries that (i) have a rating of at least one star, (ii) are single nucleotide variants, and (iii) are mapped to the Genome Reference Consortium Human Build 38 (GRCh38), using the scripts available at github.com/KULL-Centre/PRISM/tree/main/software/make prism files, release-tag v0.1.1. Next, we retained only ClinVar missense variants within the selected IDRs and omitted all deletion variants. All “likely benign” and “likely pathogenic” variants were reclassified as “benign” and “pathogenic”, respectively. Subsequently, variants with categorical labels other than “benign” and “pathogenic” were discarded.

## Data and code availability

The 29,998 conformational ensembles and precalculated conformational properties are available via sid.erda.dk/sharelink/AVZAJvJnCO which also provides access to code and data to reproduce the results presented in this work, and to `Supporting File 1.xlsx`, which lists the sequences and various sequence- and conformational properties of all 29,998 IDRs. Code and data are also available via github.com/KULL-Centre/ 2023 Tesei IDRome. CALVADOS is available from github.com/KULL-Centre/CALVADOS and can be run using Google Colab via github.com/KULL-Centre/EnsembleLab.

## Acknowledgments

We thank Drs. Tanja Mittag, Xavier Salvatella and Joshua Riback for useful comments on our work, and Dr. Alex Holehouse for sharing work before publication. We acknowledge access to computational resources from the Biocomputing Core Facility at the Department of Biology, University of Copenhagen. This work is a contribution from the PRISM (Protein Interactions and Stability in Medicine and Genomics) centre funded by the Novo Nordisk Foundation (to K.L.-L.; NNF18OC0033950). This project has received funding from the European Union’s Horizon 2020 research and innovation programme under the Marie Skłodowska-Curie grant agreement No 101025063.

## Supporting Figures

**Figure S1:**
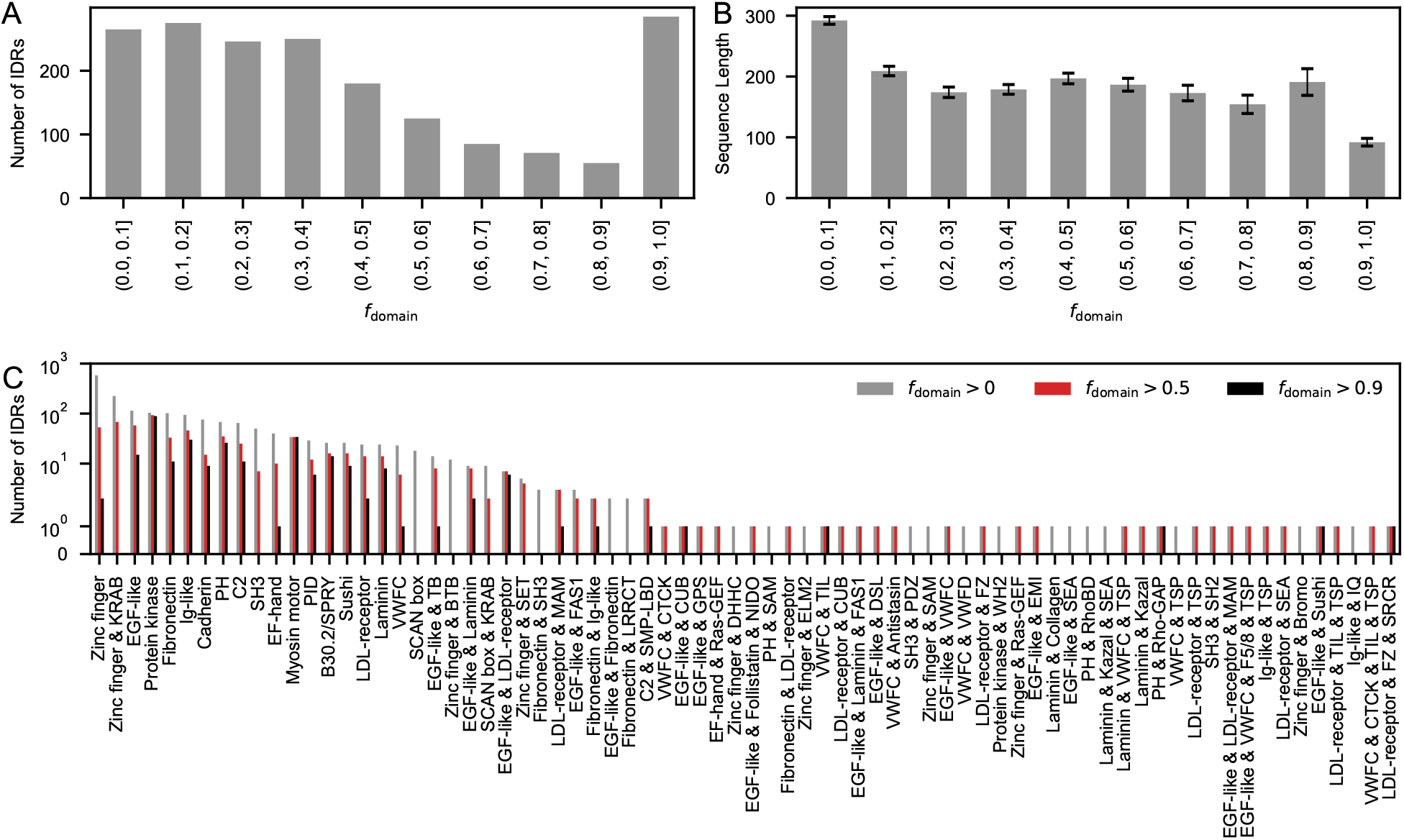
Annotation of the 1,837 IDRs (6%) with any residues in annotated domains. (*A*) Number of IDRs and (*B*) average sequence length of IDRs as a function of the fraction of residues in domains, *f*_domain_. Error bars show standard errors. (*C*) Number of IDRs in which *f*_domain_ > 0 (gray), > 0.5 (red), and > 0.9 (black) based on domain and zinc finger annotations in UniProt.

**Figure S2:**
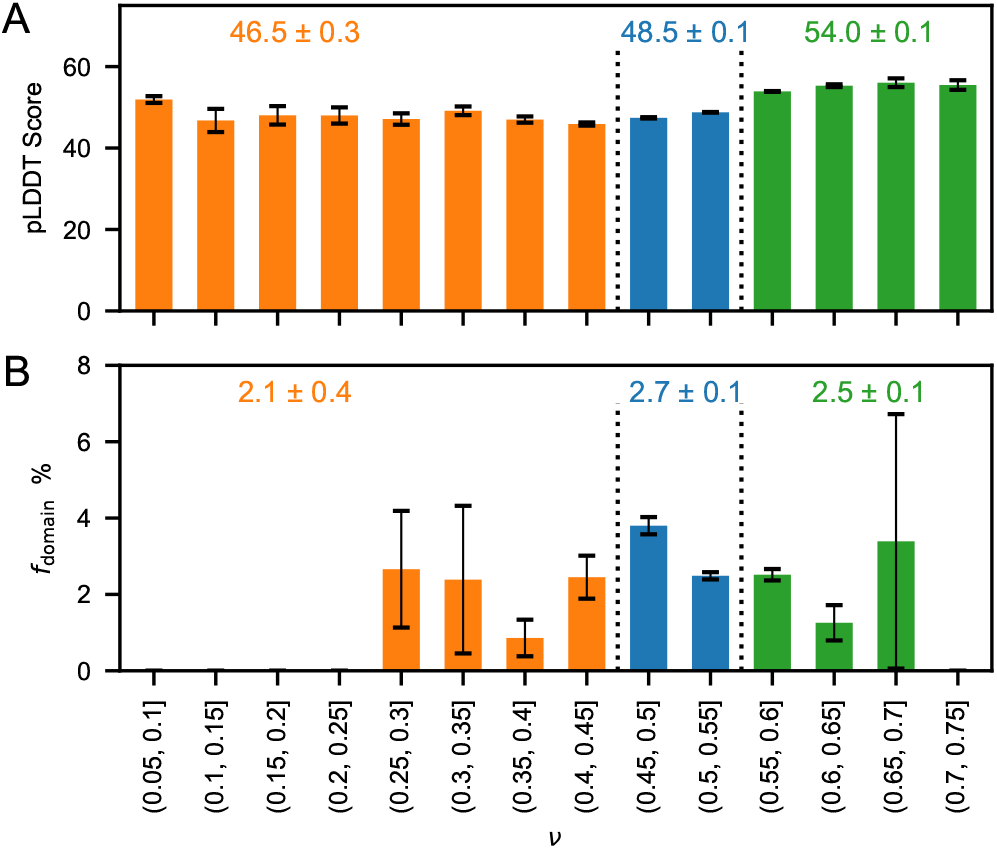
(*A*) Average pLDDT scores of the human IDRs selected in this study as a function of *ν*. (*B*) Average fraction of residues in domains annotated in UniProt for the IDRs as a function of *ν*. Error bars show standard errors and the numbers are averages and standard errors over IDRs with *ν* ≤0.45 (orange), 0.45 < *ν* ≤0.55 (blue), and *ν* > 0.55 (green).

**Figure S3:**
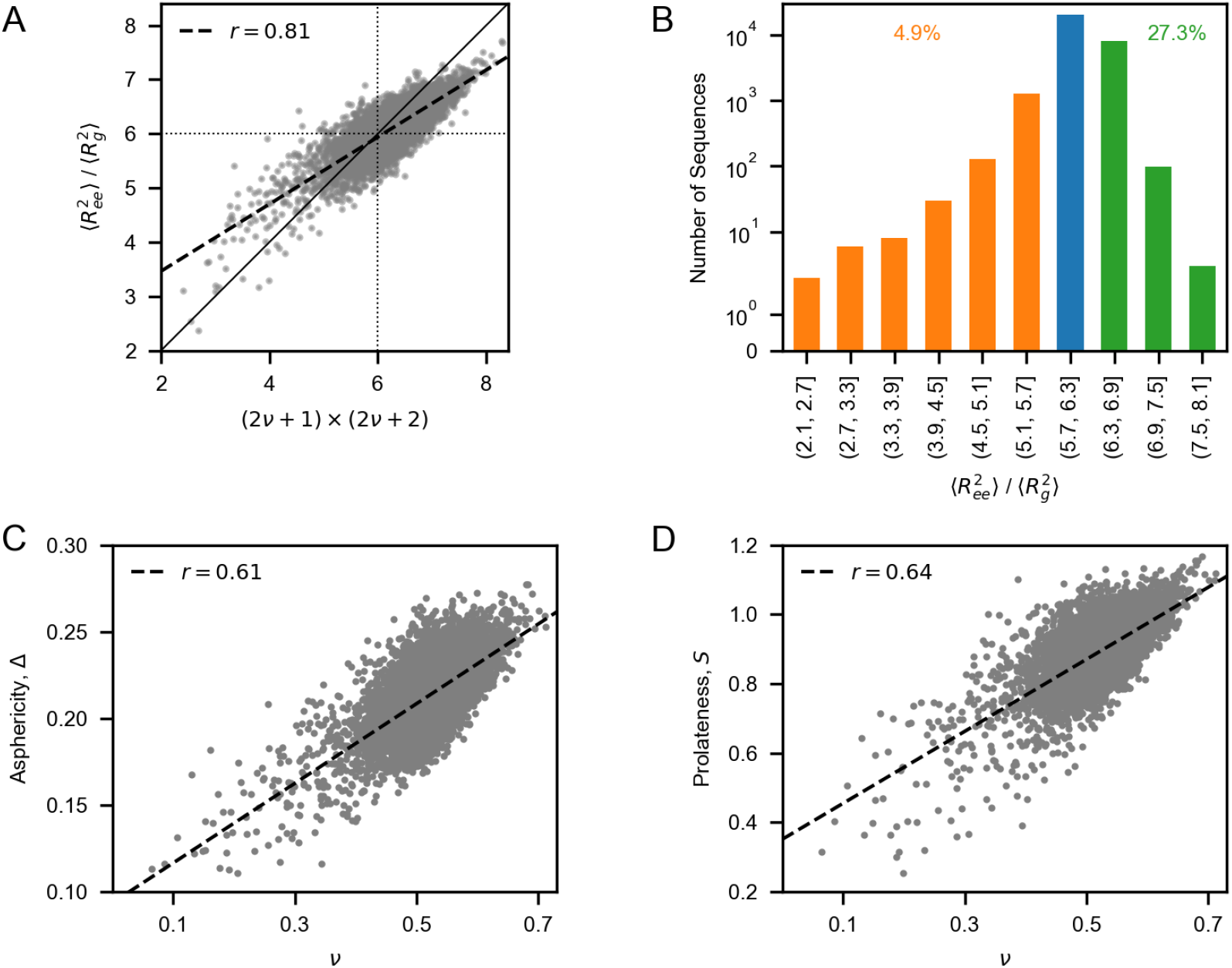
(*A*) Correlation between 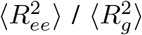 calculated from simulation trajectories and the approximate relation 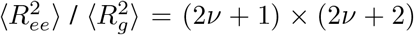, where *ν* is the apparent Flory scaling exponent [87]. (*B*) Distribution of 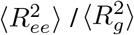 for IDRs in the human proteome; note the logarithmic scale. (*C*) Correlation between asphericity, Δ, and *ν* [88]. Correlation between prolateness, *S*, and *ν* [88]. *r* is the Pearson correlation coefficient.

**Figure S4:**
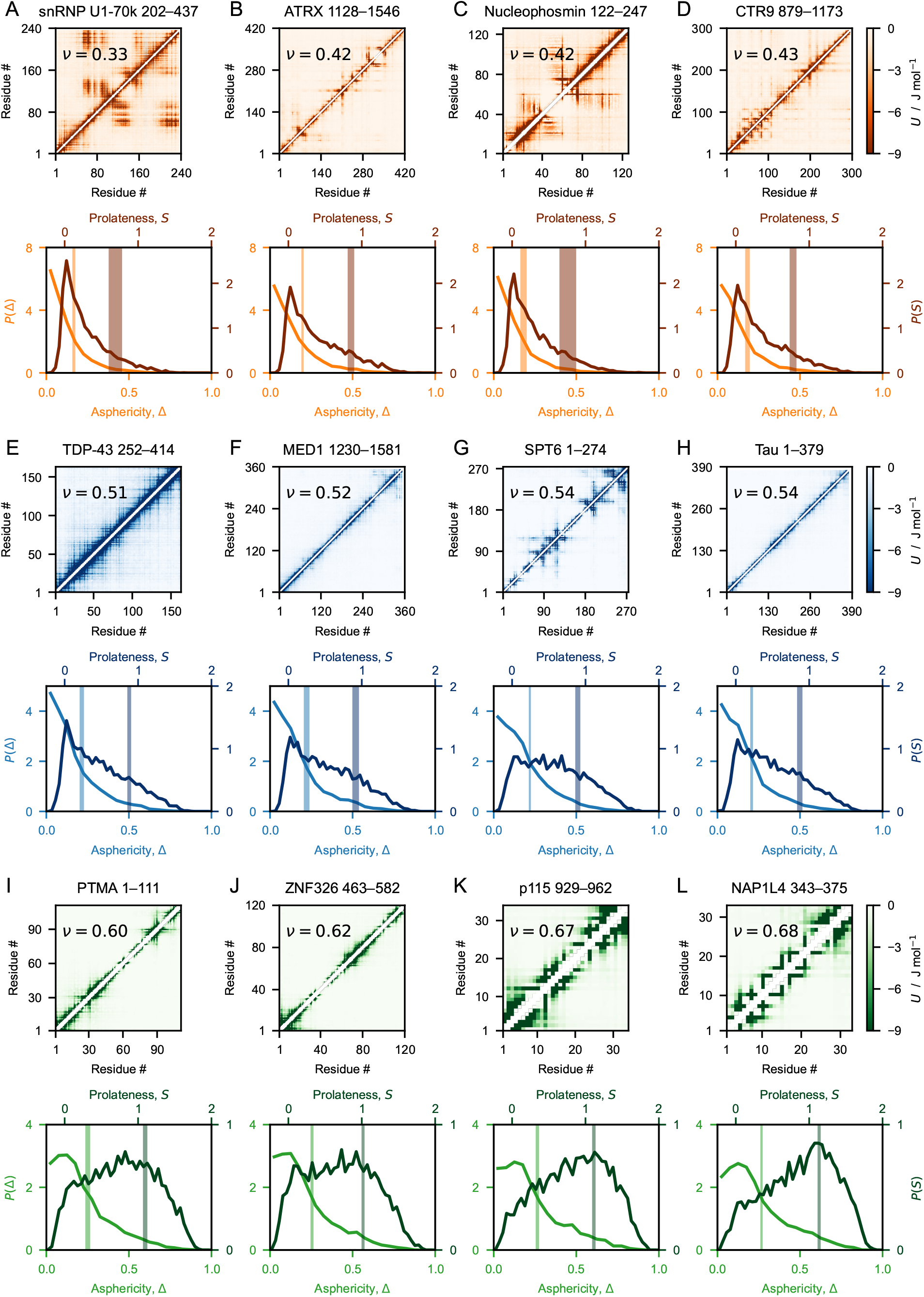
Representative intra-chain energy maps (*Top* panels), and distributions of the conformational parameters asphericity, Δ, and prolateness, *S*, (*Bottom* panels) for a selection of IDRs from our database. The UniProt IDs of the IDRs are (*A*) P08621, (*B*) P46100, (*C*) P06748, (*D*) Q6PD62, (*E*) Q13148, (*F*) Q15648, (*G*) Q7KZ85, (*H*) P10636, (*I*) P06454, (*J*) Q5BKZ1, (*K*) O60763, and (*L*) Q99733. The data is averaged over five ensembles of 1,000 weakly correlated conformations (Fig. S10), each obtained from an independent MD simulation. (*Top* panels) Energy maps are calculated using the non-ionic Ashbaugh-Hatch potential of the CALVADOS 2 model. (*Bottom* panels) Lines and vertical shaded areas show distributions and confidence intervals of the ensemble averages estimated from the standard deviation of Δ (light shade) and *S* (dark shade) over five independent simulations.

**Figure S5:**
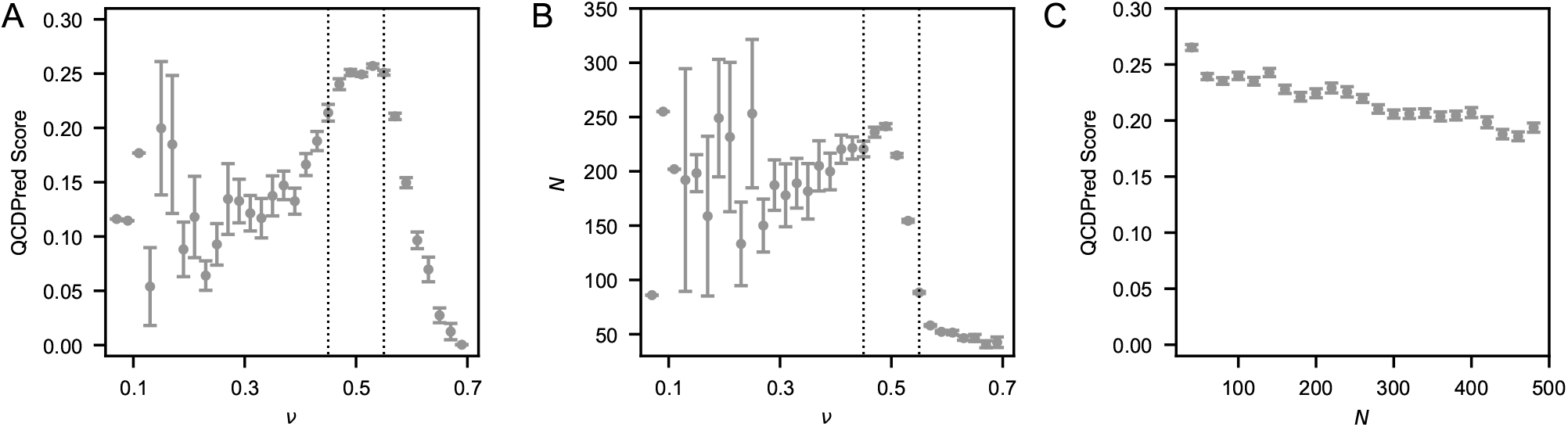
(*A*) QCDPred score and (*B*) sequence length, *N*, as a function of the apparent Flory scaling exponent, *ν*. (*C*) QCDPred score as a function of sequence length. Error bars show standard errors.

**Figure S6:**
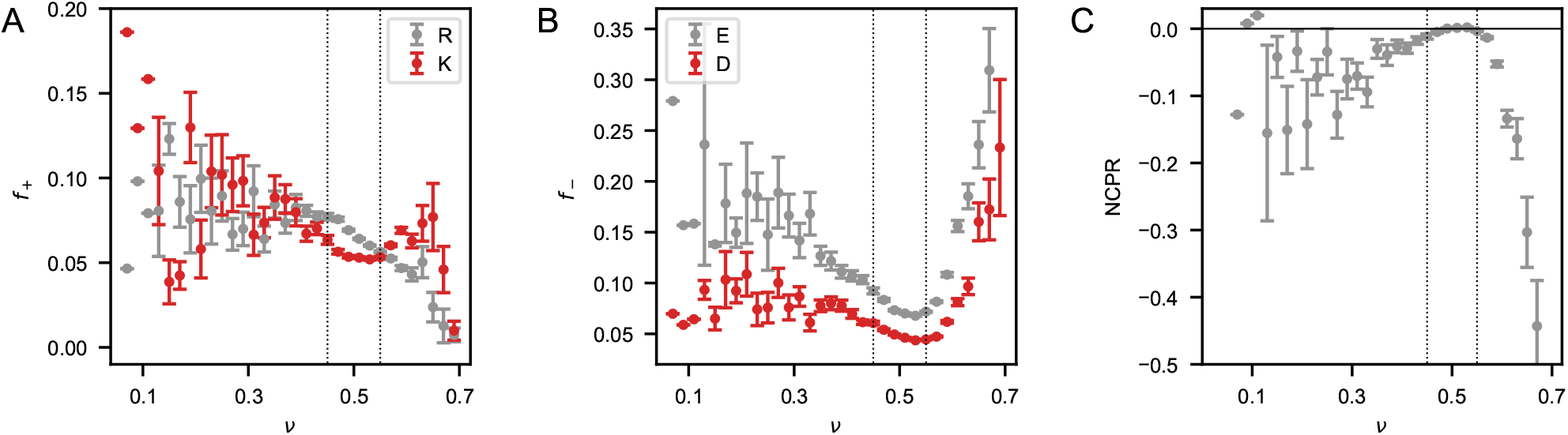
(*A*) Average fraction of positively charged residues Arg (gray) and Lys (red) as a function of *ν*. (*B*) Average fraction of negatively charged residues Glu (gray) and Asp (red) as a function of *ν*. Error bars show standard errors. (*C*) Average net charge per residue, NCPR, as a function of the apparent Flory scaling exponent, *ν*.

**Figure S7:**
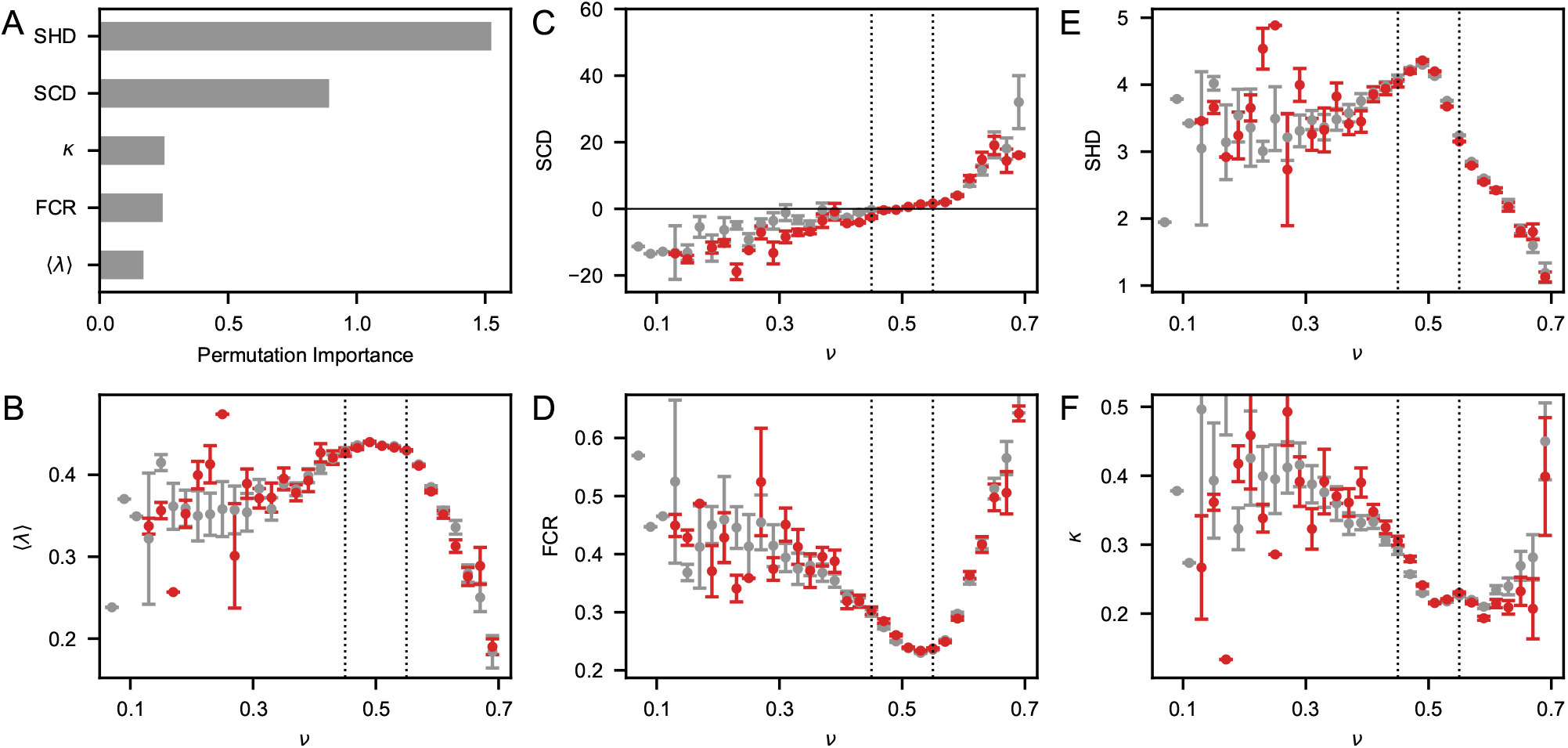
(*A*) Permutation importance of the sequence features used in the SVR model. (*B*–*F*) Average sequence features as a function of the apparent Flory scaling exponents, *ν*, calculated from simulation trajectories (gray) and using the SVR model (red). Error bars show standard errors.

**Figure S8:**
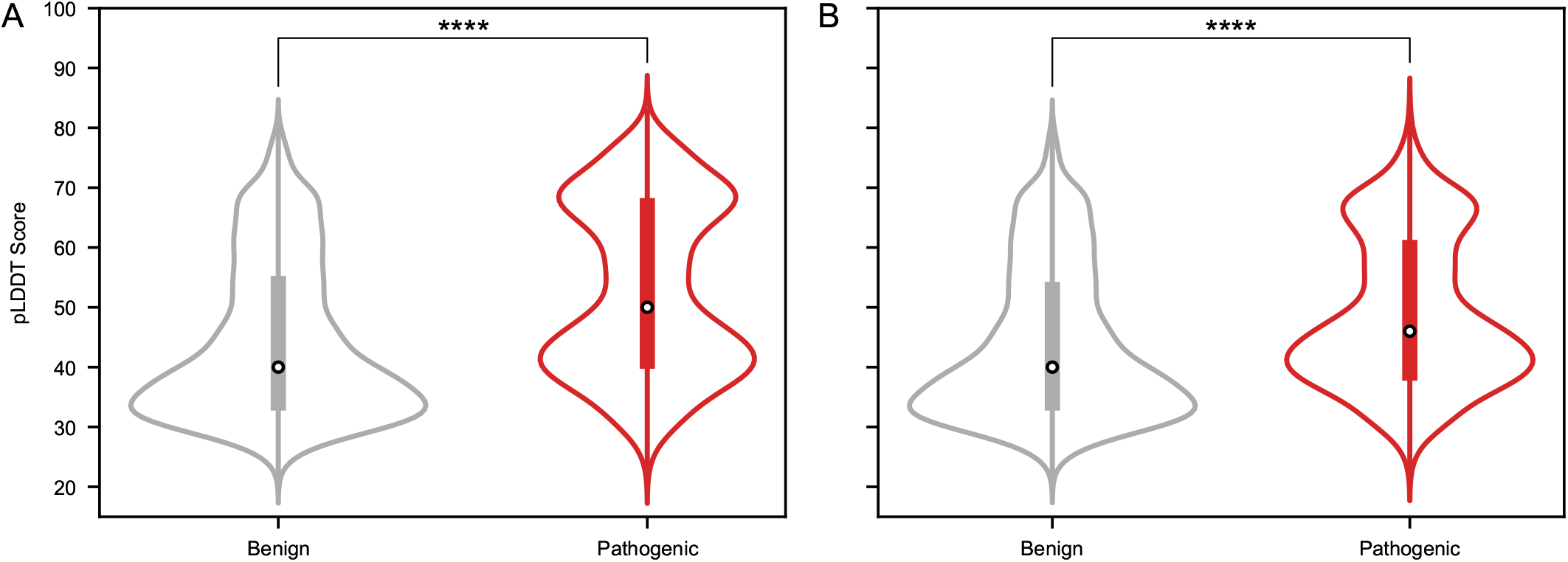
Violin plots of pLDDT scores for the wild-type residues substituted in benign (gray) and pathogenic (red) missense variants from ClinVar. The analysis is performed over variants within (*A*) all the IDRs in our database and (*B*) excluding variants within IDRs with *f*_domain_ > 0. ^****^ indicates p-values < 10^*−*75^ from a two-tailed Welch’s t-test.

**Figure S9:**
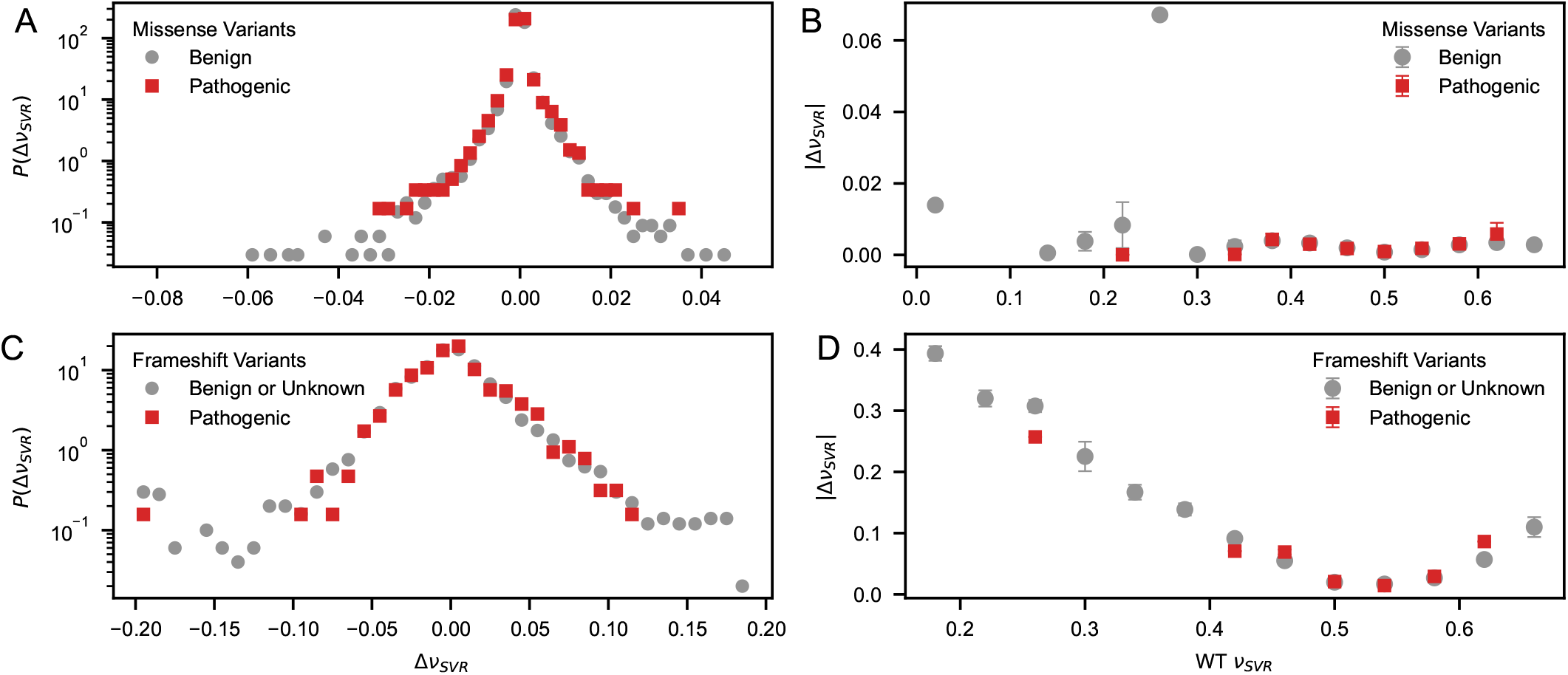
Distributions of the difference in *ν* between variant and wild-type, Δ*ν*_SVR_, for (*A*) Clinvar missense variants and (*C*) frameshift variants identified by Mensah et al. [70]. Average absolute values of Δ*ν*_SVR_ as a function of the wild-type *ν*_SVR_ value for (*B*) Clinvar missense variants and (*D*) frameshift variants identified by Mensah et al. [70]. Benign and pathogenic variants are shown in gray and red, respectively. *ν*_SVR_ values are estimated using the SVR model developed in this work.

**Figure S10:**
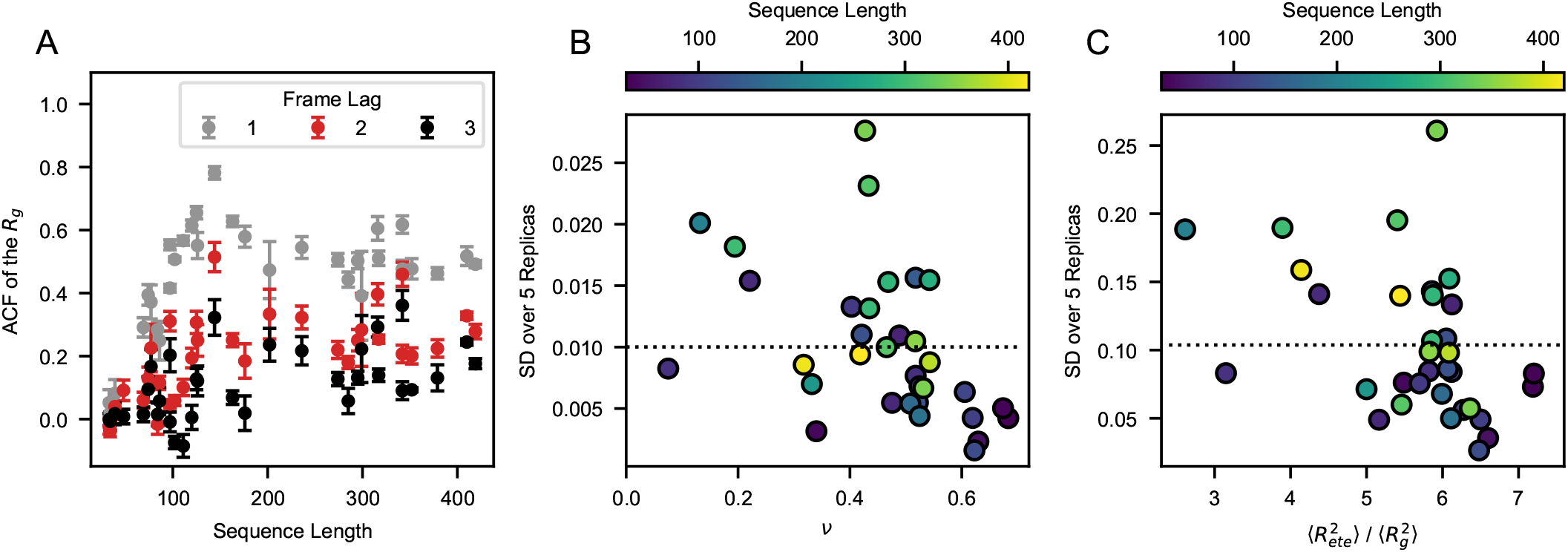
(*A*) Values of the autocorrelation function (ACF) of the *R*_*g*_ for lag times of one (gray), two (red), and three (black) frames as a function of sequence length. Error bars show standard deviation (SD) over five replicas. (*B*) Standard deviation (SD) of the apparent Flory scaling exponent, *ν*, as a function of *ν* for proteins of different sequence length. (*C*) SD of the ratio of the mean-squared end-to-end distance and the mean-squared radius of gyration, 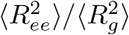, *ν*, as a function of 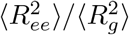 for proteins of different sequence length. SD values are calculated over five simulation replicas. Dotted lines show the average SD.

